# A Standardized Methodology for FAIRness Assessment and Multi-Dimensional Scoring in Agrosystem Research Data Infrastructures

**DOI:** 10.64898/2026.07.17.738915

**Authors:** Ata Ul Haleem, Daniel Arend, Jagadeeshwar Reddy Etukala, Elena Rey Mazón, Marcus Schmidt, Jascha Jung, Daniel Martini, Björn Usadel, Charlotte Neidiger, Robert Ulrich, Matthias Lange

## Abstract

The NFDI-consortium FAIRagro has established a systematic framework for evaluating the FAIRness of Research Data Infrastructures (RDI) within the German agrosystem research landscape. While FAIR principles are widely accepted, their practical implementation by RDIs remains challenging. By operationalizing the FAIR principles into a reproducible multi-dimensional scoring methodology, this initiative addresses the critical need for a transparent and citable benchmark of RDIs that moves beyond simple compliance. This paper details the underlying assessment criteria, comprising 20 core metrics, the iterative community-driven validation process, and the integration of these metrics into the FAIRagro Search Hub. This framework evaluates RDIs, like repositories or databases, instead of sampling hosted data sets, across the four distinct categories of FAIR independently, yielding granular, pillar-specific ratings. Unlike aggregate scoring models, which can inadvertently mask technical deficiencies by averaging performance across categories, this multi-dimensional approach ensures that a repository’s distinct strengths and bottlenecks remain fully visible. Our findings demonstrate that standardized scoring not only clarifies data accessibility for users but also highlights specific operational gaps, allowing repository providers to identify precisely where the service implementation can be enhanced. By establishing this data-driven service in the agronomy domain, we provide a scalable template for the broader NFDI and EOSC ecosystems to foster a culture of excellence in research data stewardship.

## Introduction

In the era of big data, the Findability, Accessibility, Interoperability, and Reusability (FAIR) of datasets are no longer aspirational but are foundational requirements for scientific progress and the mitigation of the reproducibility crisis (Wilkinson et al., 2016). While the FAIR principles provide a high-level theoretical framework, translating these abstract guidelines into measurable, domain-specific metrics remains a significant technical challenge (Mons et al., 2017). Many repositories have a long history and have operated well for their communities to preserve data but fall short in some cases in serving as part of a federation of Research Data Infrastructures due to the lack of standardized metadata or interoperable interfaces (Maemura, 2021).

Within the German National Research Data Infrastructure (NFDI), a federated system of decentralized RDIs is critical for achieving cross-disciplinary data integration (Diepenbroek et al., 2014). However, a significant gap remains: while federated RDIs share common FAIR goals, the lack of unified technical baselines often leads to fragmented FAIRness for the hosted data sets that affects data discovery and cause inconsistent metadata across decentralized nodes (Stockhause & Lautenschlager, 2017). This is particularly evident in the multidisciplinary agrosystems domain spanning soil science, genomics, and socio-economics repositories, each with varying levels of digital maturity (Schlindwein, 2025).

To address these challenges without imposing a rigid top-down structure, the FAIRagro initiative proposes a tailored approach starting from the agrosystem landscape in Germany. By suggesting a FAIR assessment framework operationalizing the RDA FAIR Data Maturity Indicators (RDA FAIR Data Maturity Model Working Group, 2020) through a four-pillar framework, FAIRagro initiative aims to move beyond abstract theory toward a practical assessment that respects the diversity of federated infrastructures. This approach facilitates the convergence of decentralized RDIs (Curdt et al., 2022) into a multi-domain metadata space; within this framework, FAIRness is defined as the measurable, evolving capability of an RDI to systematically enable and unify Findability, Accessibility, Interoperability and Reusability across its holdings. This approach shifts the evaluation from individual data sets to the FAIR-enabling infrastructure that governs them.

A technical realization and validation of this assessment methodology are implemented via a dedicated standalone prototype for FAIRagro Repository Search (https://repofinder.ipk-gatersleben.de/), which provides an active gateway to explore these assessments. Rather than relying on the rigid, pre-computed aggregate scores generated by typical automated assessment tools (Azevedo et al., 2025; Candela et al., 2024; Devaraju & Huber, 2020; Gaignard et al., 2023), which often mask underlying infrastructural variations, the FAIR RDI dashboard exposes individual properties at a baseline level of granularity. This design empowers researchers to execute user-driven queries and prioritize repository features based on their specific project requirements and workflows. In contrast to prescriptive computational rankings, this study presents a robust methodology to discover relevant RDIs through flexible, user-defined feature profiles, providing a citable reference that aligns with the NFDI’s strategic mandate (Diepenbroek et al., 2023; Zänkert et al., 2025) for transparent and interoperable research data management.

## Methods

The assessment framework is operationalized through three distinct methodological components:

### Metric Definition

The FAIRagro assessment framework operationalizes the RDA FAIR Data Maturity Indicators (RDA FAIR Data Maturity Model Working Group, 2020) into a reproducible, multi-dimensional scoring methodology tailored to the German agrosystem research landscape. The framework evaluates infrastructures across twenty core FAIR metrics, comprising four pillars of five distinct binary attributes (1 = compliance, 0 = non-compliance), yielding a granular, non-aggregated score for each principle. The criteria for each category are illustrated in Figure 1.

**Figure 1:**
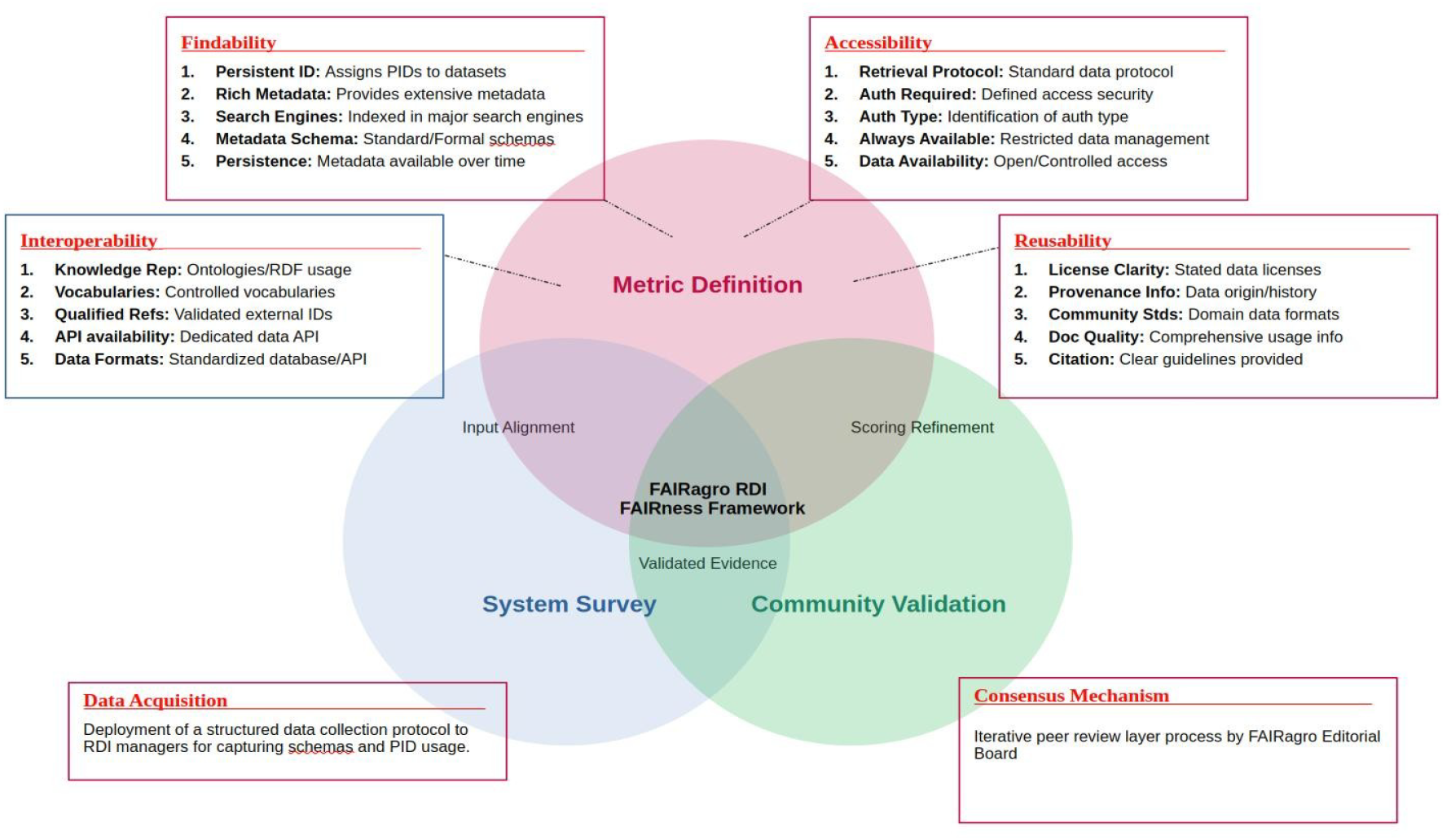
The FAIRagro RDI FAIRness Framework Methodology. The assessment framework is built upon three intersecting pillars: Metric Definition (Top) that establishes the 20 core criteria across Findability, Accessibility, Interoperability, and Reusability (FAIR), ranging from persistent identifier assignment to community metadata standards, the System Survey (Bottom Left) representing the data acquisition phase, where structured protocols are deployed to RDI managers to capture technical schemas and evidence and the Validation (Bottom Right), a consensus mechanism involving iterative peer-review within FAIRagro Editorial Board to refine scoring and ensure domain relevance. The intersections represent critical alignment phases: *Input Alignment* (translating survey data to metrics), *Scoring Refinement* (validating metrics against community needs), and *Validated Evidence* (the final synthesis of empirical data and expert consensus).

### Data Sources and Acquisition

To systematically populate the FAIRness property catalogue, a hybrid data acquisition pipeline was developed (https://github.com/fairagro/rdi-fairness-interviews) that curates repository FAIRness data from two complementary sources 1) an interview as questionnaire to the RDI operators and, if one exists, 2) RDIs self-contributed, community-reviewed profiles to re3data, further complementing 3) with reviews and curation by FAIRagro data stewards.

To this end, a structured questionnaire based on the modified RDA framework was distributed to the responsible managers of every RDI associated with the FAIRagro consortium. This enabled direct, objective reports that provided expert-level insight into the RDI implementation. Crucially, they supply explicit evidence for features that are guessed by heuristics implemented in FAIR-assessment tools or inferred from global RDI registries only. Examples are quality of user documentation, internal provenance tracking configurations, and long-term metadata persistence guarantees, etc. The complete interview profiles and questionnaire templates are available for download (https://repofinder.ipk-gatersleben.de/downloads). For those RDIs where the questionnaire is pending, the automated harvesting pipeline (https://github.com/fairagro/rdi-fairness-interviews/tree/main/rdiDataEnricher) was implemented to extract repository characteristics from the re3data.org registry API, if a re3data record exists. The pipeline translates raw registry elements and RDI interview responses into the twenty standardized FAIR indicators. To this end, the re3data attributes are strictly compartmentalized into three distinct provenance categories to preserve data lineage, native (Attributes extracted via direct, 1:1 mapping from explicit re3data XML elements), inferred (Attributes requiring semantic or text-based heuristics to determine compliance) and novel (Highly granular technical milestones entirely unrepresented within the standard re3data metadata schema). (e.g., Metadata Persistence Policies, Formal Knowledge Representation/Ontologies, and Dataset Provenance Information). These attributes can only be populated via direct operator interviews or manual curation by domain data stewards.

### Community Validation Loop

To ensure the actual accuracy of both automated inferences and self-reported metrics, all compiled profiles undergo an iterative validation loop. Preliminary scores are subjected to a peer review by a panel of domain data stewards within the FAIRagro Data Steward Service Center (DSSC) Editorial Board. The features of the final version are pushed directly to the public web UI (https://repofinder.ipk-gatersleben.de/downloads) and made available through api request (e.g. https://repofinder.ipk-gatersleben.de/api/resource/repository/RFId009202604272) as well as a publicly available npm package (https://www.npmjs.com/package/rf-rdis) for integration in data curation workflows.

## Results

This section presents the results of the FAIRness assessment survey conducted across RDIs in the agrosystem research domain. The analysis is based on the standardized methodology described previously, using twenty core FAIR metrics grouped under Findability, Accessibility, Interoperability, and Reusability derived from RDA maturity metrics.

### Aggregate FAIR Maturity of FAIRagro Agrosystem RDIs

To evaluate the overall digital maturity of the agrosystem repository landscape while exposing systematic gaps in registry-level tracking frameworks, we analyzed the compliance profiles across three distinct analytical configurations: (A) a high-accuracy, hand-curated interview group of re3data repository operators (n=14); (B) an isolated, automated fallback population also harvested from re3data (n=31) and mapped to FAIR attributes; and (C) an overall landscape baseline containing both A and B. The separate, pillar-specific averages for these three dimensions are overlaid on a standardized 0–5 scale grid in Figure 2. As illustrated in the radar chart, the hand-curated survey repositories show high maturity in Reusability, Findability and Interoperability (averaging 4.64, 4.57 and 4.07 out of 5, respectively), with Accessibility lower at 3.5, largely because few open repositories require authentication.

**Figure 2:**
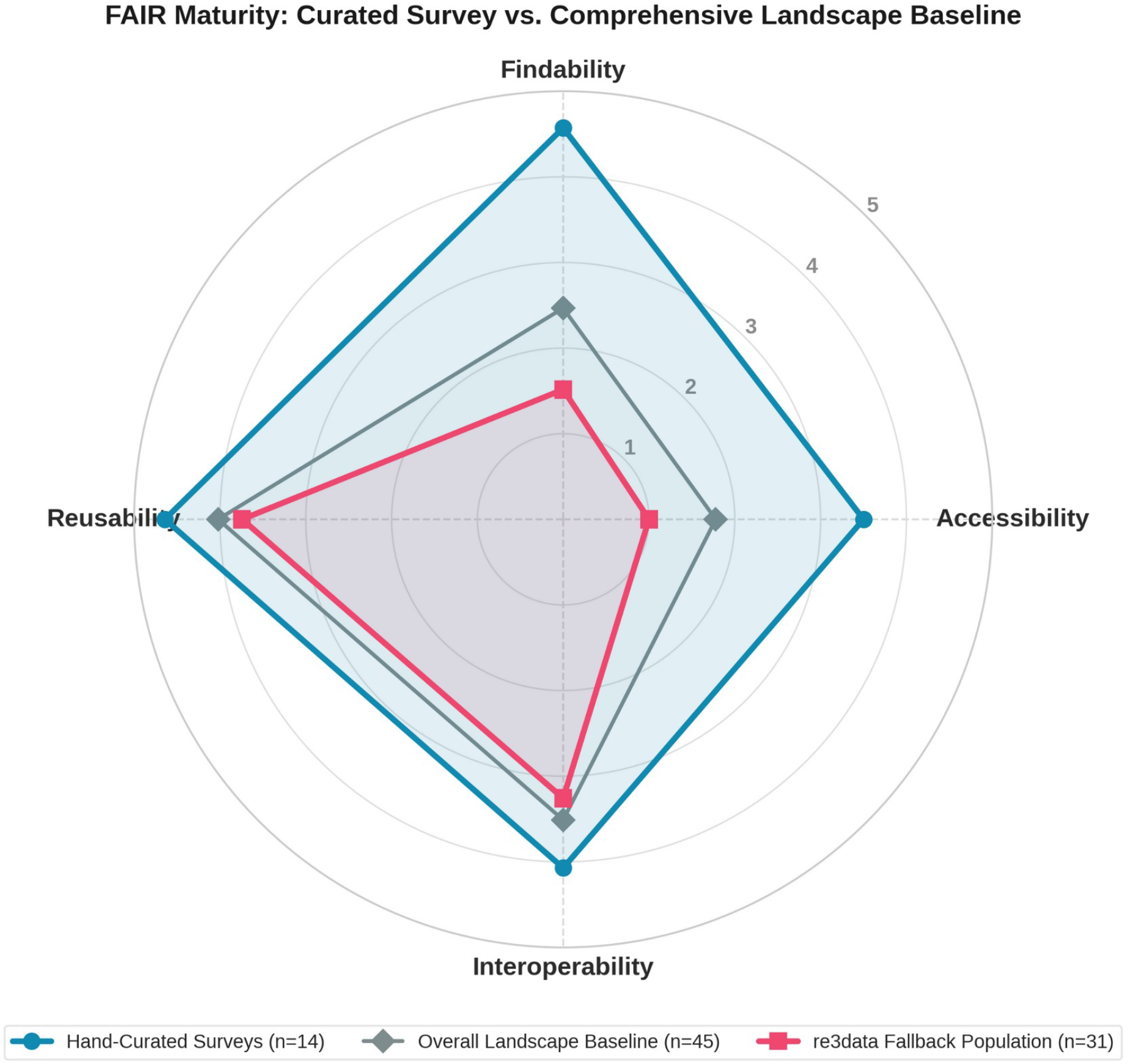
Multi-tier comparative analysis of FAIR maturity across the FAIRagro agrosystem repository landscape. The radar chart illustrates average compliance scores across the four fundamental pillars of the FAIR principles on a standardized 0–5 scale. Three distinct analytical dimensions are mapped to evaluate infrastructure digital maturity and assess the visibility of automated indexing: Hand-Curated Surveys (blue circle), re3data Fallback Population (pink square) and Overall Landscape Baseline (gray diamond)

In comparison, the isolated re3data fallback population displays a highly focused profile that reflects its intended design as a high-level, cross-disciplinary discovery index. This cohort reaches its highest marks in Reusability (approximately 3.74 out of 5) and Interoperability (3.26 out of 5), while maintaining a foundational baseline in Findability (1.52 out of 5) and Accessibility (1.0 out of 5). By integrating these configurations into a single composite view, the resulting global landscape baseline settles at an intermediate equilibrium, tracking at approximately 3.5 for Interoperability, 1.78 for Accessibility, 2.47 for Findability, and 4.02 for Reusability.

### Detailed Indicator Compliance identify key strengths and weaknesses of FAIRagro Agrosystem RDIs

The bar chart in Figure 3 provides a granular view of compliance rates for each FAIR indicator across the distinct evaluation cohorts, revealing specific technical dimensions where the landscape excels or faces structural limitations. As illustrated in Figure 3, the RDI-operator interviews (n=14) show maximum compliance (100%) across multiple fundamental features, including core criteria like licenseClarity, dataAvailability, metadataAlwaysAvailable, retrievalProtocol, metadataSchema, and richMetadata. Near-perfect compliance (≈92%) is similarly observed in fields like documentationQuality, citationGuidelines, communityStandards, dataFormats, apiAvailability, and metadataPersistence.

**Figure 3:**
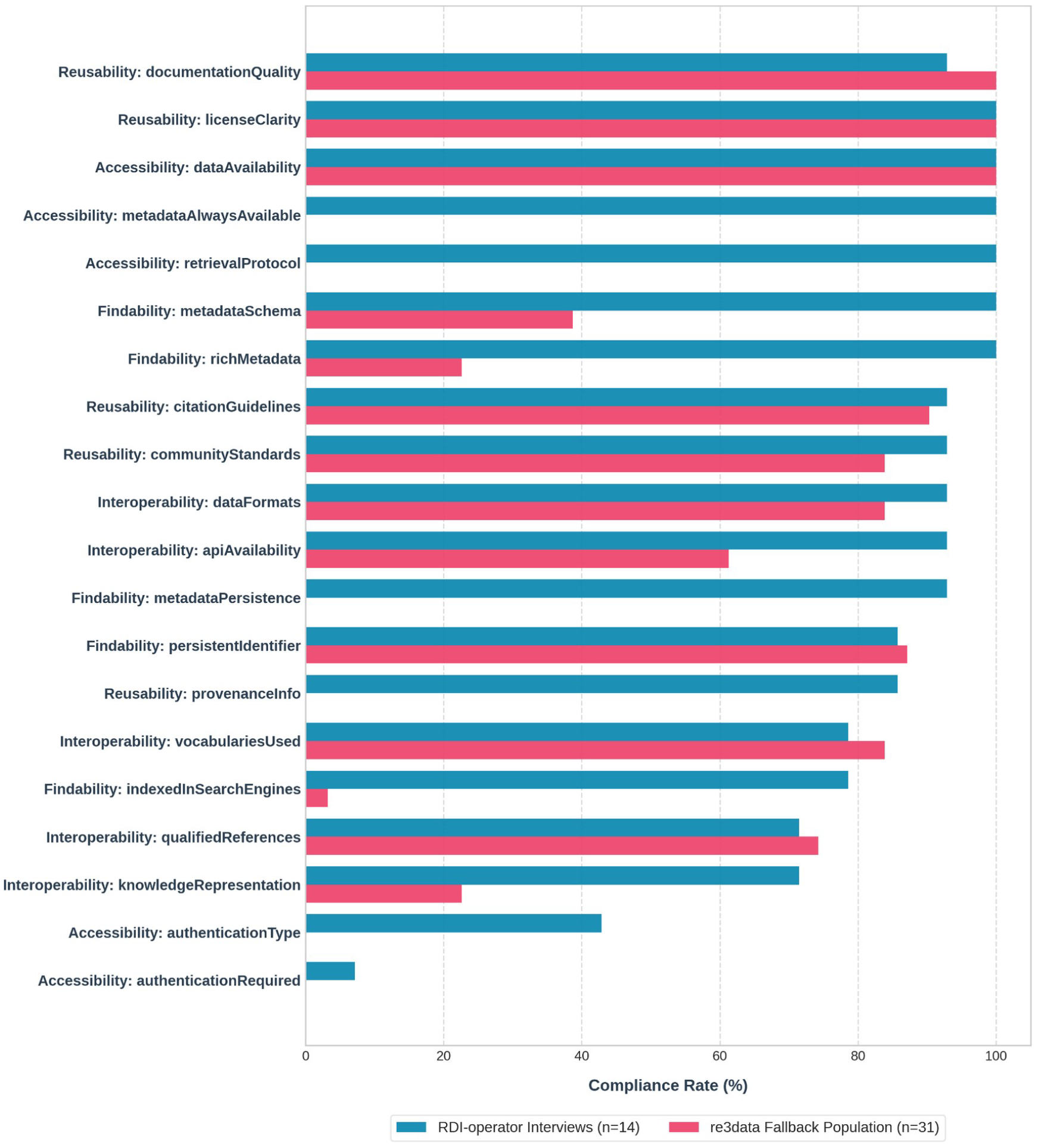
Technical indicator compliance comparison. The chart maps the percentage of repositories meeting specific FAIR criteria within the high-fidelity RDI-operator Interviews (blue, n=14) versus the automated re3data Fallback Population (pink, n=31).

When contrasted with the re3data fallback population (n=31), the variations reflect the inherent granularity limits of broad indexing schemas rather than a failure of the repositories themselves. For metrics like *metadataAlwaysAvailable*, and *retrievalProtocol*, where the interviewed repositories achieve absolute compliance, the re3data fallback group registers below 5%. The drop indicates that these specific technical executions occur via localized protocols that fall outside the standard, un-enriched re3data harvesting scope. This is supported by metrics where the two populations align or where the registry is highly effective documentationQuality yields high metrics across both cohorts (92% for interviews vs. 100% for re3data), and vocabulariesUsed actually shows strong visibility in re3data (85%) compared to the interview group (77%), indicating the registry’s efficacy at capturing standard vocabulary use.

Conversely, for other indicators, the low metrics may reflect actual structural data gaps within the wider repository ecosystem rather than a mere harvesting issue. For example, persistentIdentifier demonstrates exceptional alignment and high compliance across both cohorts (∼86% among interviewed repositories and ∼87% for the fallback group), suggesting that robust PID assignment workflows are thoroughly established throughout the broader landscape of RDIs. On the other hand, compliance for formal knowledge representation schemas (*knowledgeRepresentation*) is constrained, dropping from ∼72% for interviews to 23% for re3data, indicating that structured semantic web bindings remain a widespread technical challenge. Furthermore, the requirement for user authentication frameworks (*authenticationRequired* and *authenticationType*) displays low values or complete absence across both groups, highlighting areas where technical capabilities are genuinely restricted or unaligned across the landscape.

### Implementation: Use cases for FAIRagro RDI FAIRness Framework

The primary outcome of this methodology is the live operational deployment of the FAIRness assessment layer, transitioning the framework from a theoretical benchmarking model into a set of machine-actionable services. By moving beyond static, flat reports, the system associates multi-dimensional FAIRness metadata directly with verified repository identities through a decoupled, multi-tiered deployment architecture. This architecture is operationalized across the following core results:

#### 1. PID-Linked FAIRness Profiles (RFId URIs)

A foundational achievement is the successful assignment of unique RepoFinder time-stamped (TS) identifiers (RFIds) to 77 repositories across 27 scientific domains. These persistent identifiers (Supplementary Figure 2) act as resolvable anchors for all underlying FAIRness metadata. By cross-referencing and binding these RFIds to established global registry identifiers (such as re3data IDs), the system guarantees strict data lineage and facilitates seamless programmatic cross-referencing across federated infrastructures.

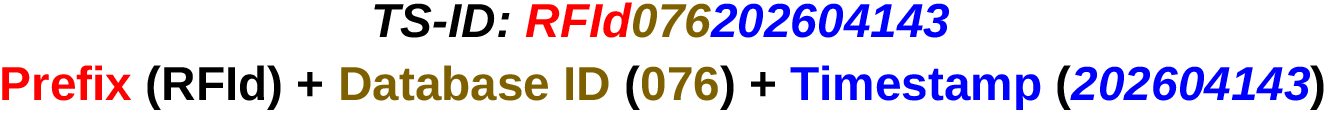

#### 2. Visual User Interface

The framework operationalizes human-readable discovery through two primary web-based visual layers:

- **The Repository Overview Card** The primary representation for repositories is the Repository Overview Card (Figure 4) using the BonaRes repository as a representative example. This component serves as a dense, high-level summary container and as a quick snapshot of the metadata associated with the corresponding RDI, its operational profile, and a high-level FAIRness evaluation. To facilitate consistent interpretation across the platform, Table 1 provides a detailed taxonomic breakdown of the symbolic attribution used to represent each of the twenty core FAIR metrics.
- **Comparative Search Integration:** To facilitate rapid, high-level benchmarking during repository search, “FAIRness cards” are embedded directly into the search results page (https://repofinder.ipk-gatersleben.de/). This interface allows users to visually compare the ratings of multiple individual FAIRness properties simultaneously before diving into comprehensive repository documentation (Supplementary Figure 1).
- **Granular Evaluation Dashboards:** By structuring assessment data as a resolvable resource (/resource/repository/{**RFId**}/fairness), the framework generates deep-dive visual profiles for each infrastructure (e.g., https://repofinder.ipk-gatersleben.de/resource/repository/RFId001202604272/ fairness). As illustrated in the BonaRes repository profile (Figure 5), granular metric compliance is synthesized into an interactive bar chart. This immediately exposes technical strengths and operational bottlenecks, signalling the roadmap for improvements.
- **Application Programming Interface (API) for real time RDI-FAIRness comparison** For large-scale metadata harvesting, automated infrastructure auditing, and cross-consortium analysis, the framework exposes a dedicated, production-ready RESTful API layer (https://repofinder.ipk-gatersleben.de/api-docs). Developers and data stewards can programmatically query the full, live RDI metadata object, including schema-compliant details on institutional provenance, supported metadata standards (e.g., Dublin Core), and use standard API calls structured around the repository’s identifier. Additionally, URIs of the PID-Linked FAIRness Profiles enable the automatic, continuous, and unambiguous monitoring of the technical nuances of the federated landscape by external validation engines (e.g., https://fairagro.github.io/rdi-fairness-interviews/api/resource/repository/RFId001202604272/fairness). Figure 6 details the machine-actionable response returned by our FAIRness API.

**Table 1:**
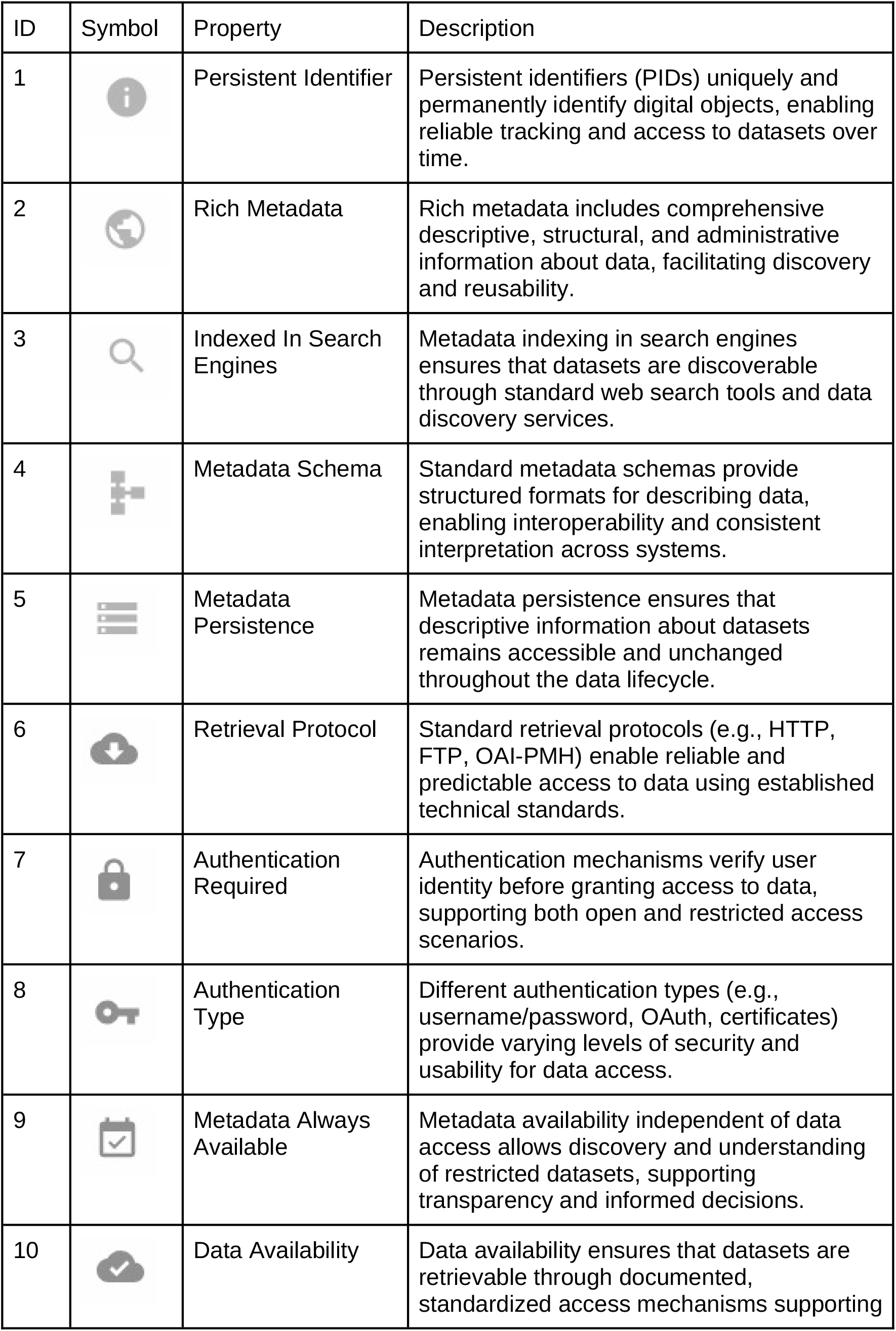
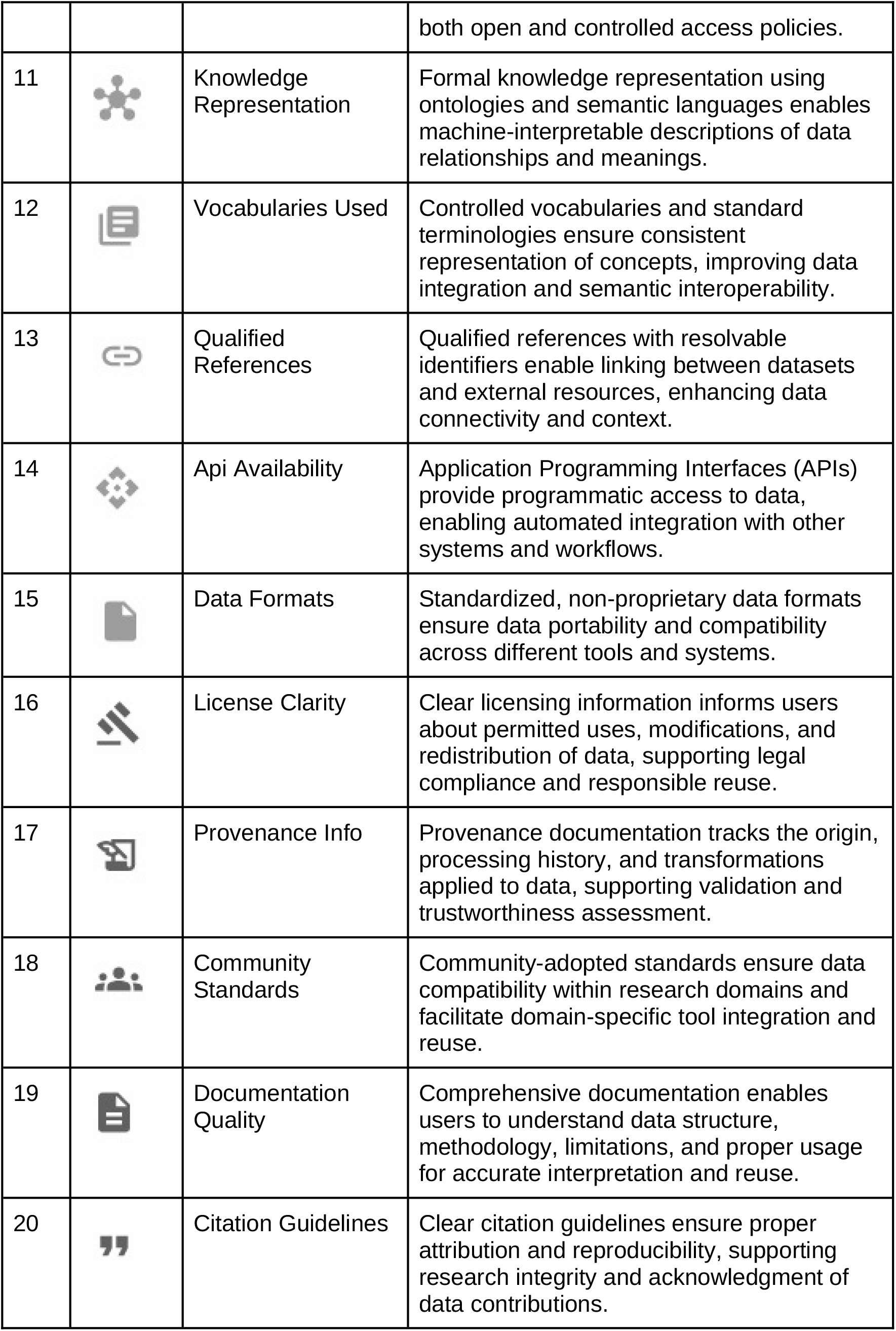
Symbolic mapping of the twenty core FAIRness evaluation criteria, including corresponding visual symbols, properties (RDA indicators) and their description.

| ID | Symbol | Property | Description |
| --- | --- | --- | --- |
| 1 |  | Persistent Identifier | Persistent identifiers (PIDs) uniquely and permanently identify digital objects, enabling reliable tracking and access to datasets over time. |
| 2 |  | Rich Metadata | Rich metadata includes comprehensive descriptive, structural, and administrative information about data, facilitating discovery and reusability. |
| 3 |  | Indexed In Search Engines | Metadata indexing in search engines ensures that datasets are discoverable through standard web search tools and data discovery services. |
| 4 |  | Metadata Schema | Standard metadata schemas provide structured formats for describing data, enabling interoperability and consistent interpretation across systems. |
| 5 |  | Metadata Persistence | Metadata persistence ensures that descriptive information about datasets remains accessible and unchanged throughout the data lifecycle. |
| 6 |  | Retrieval Protocol | Standard retrieval protocols (e.g., HTTP, FTP, OAI-PMH) enable reliable and predictable access to data using established technical standards. |
| 7 |  | Authentication Required | Authentication mechanisms verify user identity before granting access to data, supporting both open and restricted access scenarios. |
| 8 |  | Authentication Type | Different authentication types (e.g., username/password, OAuth, certificates) provide varying levels of security and usability for data access. |
| 9 |  | Metadata Always Available | Metadata availability independent of data access allows discovery and understanding of restricted datasets, supporting transparency and informed decisions. |
| 10 |  | Data Availability | Data availability ensures that datasets are retrievable through documented, standardized access mechanisms supporting both open and controlled access policies. |
| 11 |  | Knowledge Representation | Formal knowledge representation using ontologies and semantic languages enables machine-interpretable descriptions of data relationships and meanings. |
| 12 |  | Vocabularies Used | Controlled vocabularies and standard terminologies ensure consistent representation of concepts, improving data integration and semantic interoperability. |
| 13 |  | Qualified References | Qualified references with resolvable identifiers enable linking between datasets and external resources, enhancing data connectivity and context. |
| 14 |  | Api Availability | Application Programming Interfaces (APIs) provide programmatic access to data, enabling automated integration with other systems and workflows. |
| 15 |  | Data Formats | Standardized, non-proprietary data formats ensure data portability and compatibility across different tools and systems. |
| 16 |  | License Clarity | Clear licensing information informs users about permitted uses, modifications, and redistribution of data, supporting legal compliance and responsible reuse. |
| 17 |  | Provenance Info | Provenance documentation tracks the origin, processing history, and transformations applied to data, supporting validation and trustworthiness assessment. |
| 18 |  | Community Standards | Community-adopted standards ensure data compatibility within research domains and facilitate domain-specific tool integration and reuse. |
| 19 |  | Documentation Quality | Comprehensive documentation enables users to understand data structure, methodology, limitations, and proper usage for accurate interpretation and reuse. |
| 20 |  | Citation Guidelines | Clear citation guidelines ensure proper attribution and reproducibility, supporting research integrity and acknowledgment of data contributions. |

**Table 2:** API reference for rf-rdis, npm package.

| Function call | Action |
| --- | --- |
| `getAllRdis()` | Retrieves the full, unified array of repository metadata objects registered in the system. |
| `getRdiById(id)` | Fetches a single repository's metadata envelope using its unique persistent identifier. |
| `searchRdis(predicate)` | Enables client-side predicate-based searches, allowing developers to filter repositories using custom parameters (e.g., selecting repositories containing domain keywords like "Soil"). |
| `getFairnessById(id)` | Directly extracts the schema-compliant `fairnessAssessment` object mapped to the specified identifier. |

**Figure 4:**
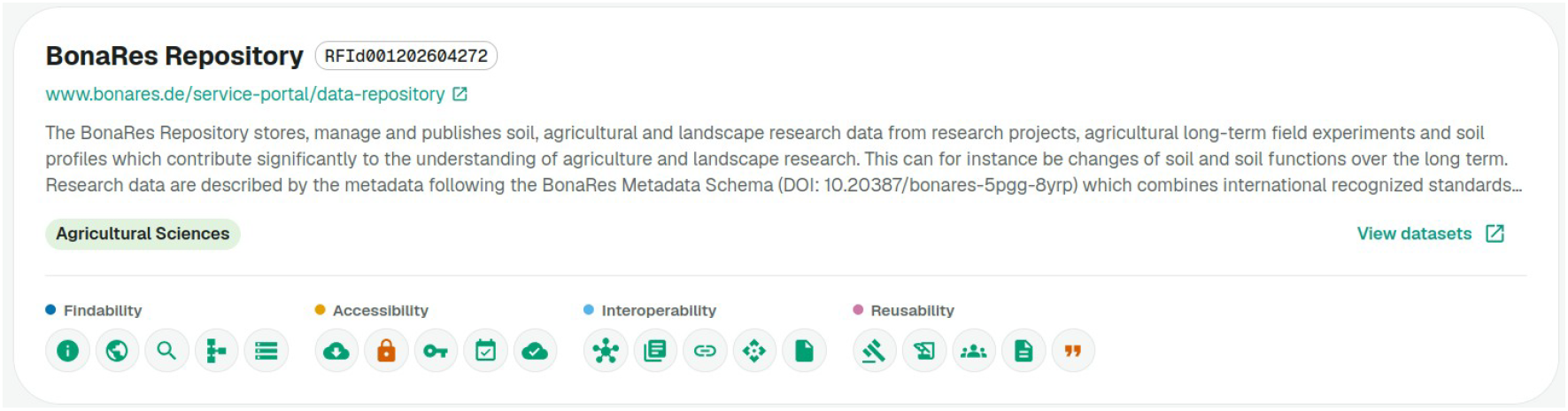
User interface layout of the Repository Overview Card showing registry metadata and the inline 20-indicator FAIRness diagnostic grid. The card is split into three core functional areas: (1) Identity & Location Metadata at the top, displaying the repository name (BonaRes Repository), its FAIRagro specific persistent unique identifier (RFId001202604272), and direct access URL; (2) Repository Context in the middle, including a detailed text description of its domain focus; and (3) the FAIRness Diagnostic Grid at the bottom. This diagnostic footer uses a visual traffic-light color-coding system to depict compliance across 20 distinct FAIR metrics, where green icons denote full compliance with the metric, red indicates non-compliance, and grey denotes an unknown status.

**Figure 5:**
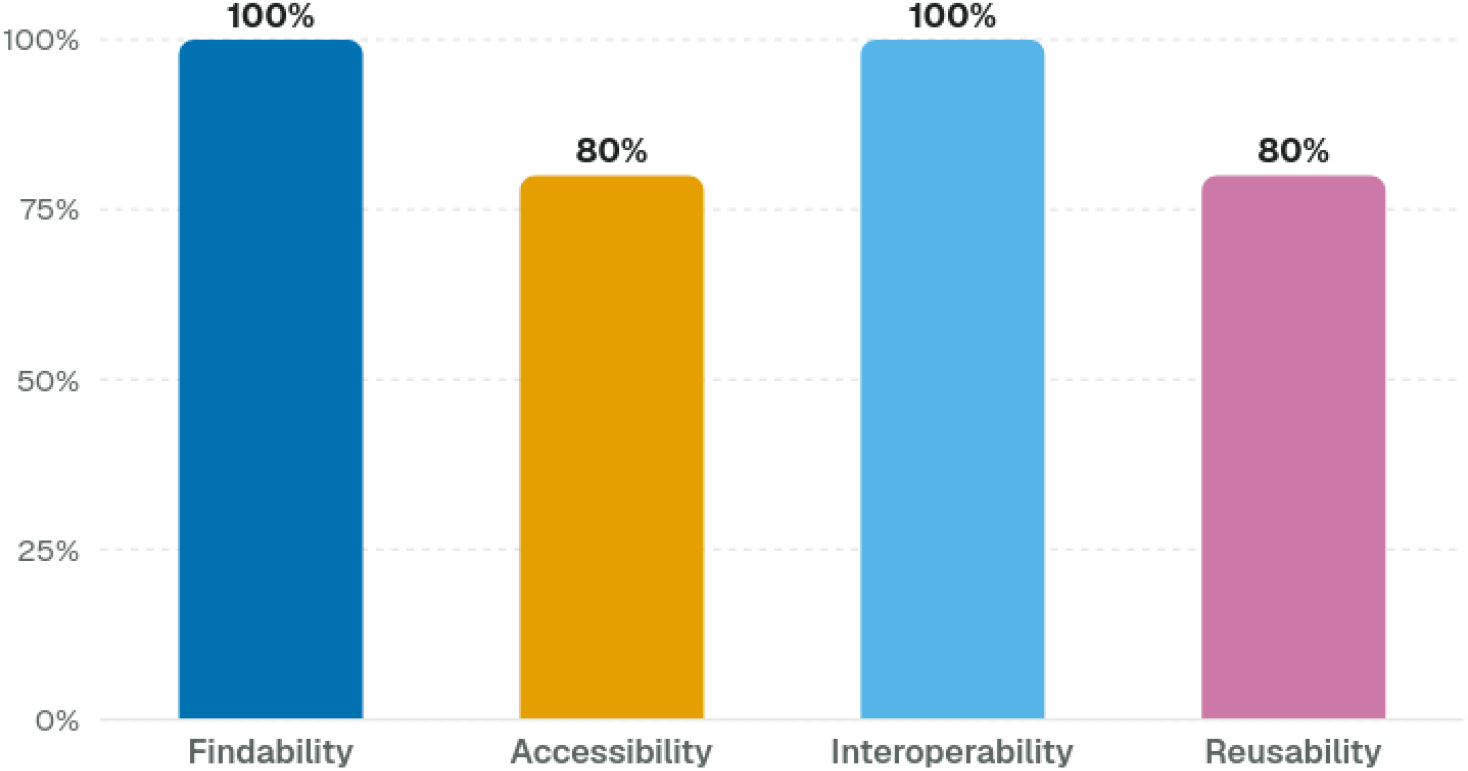
FAIRness Assessment Results for the Bonares Repository. The bar chart illustrates the granular scoring as percentage across the four FAIR pillars on a 5-point scale.

**Figure 6:**
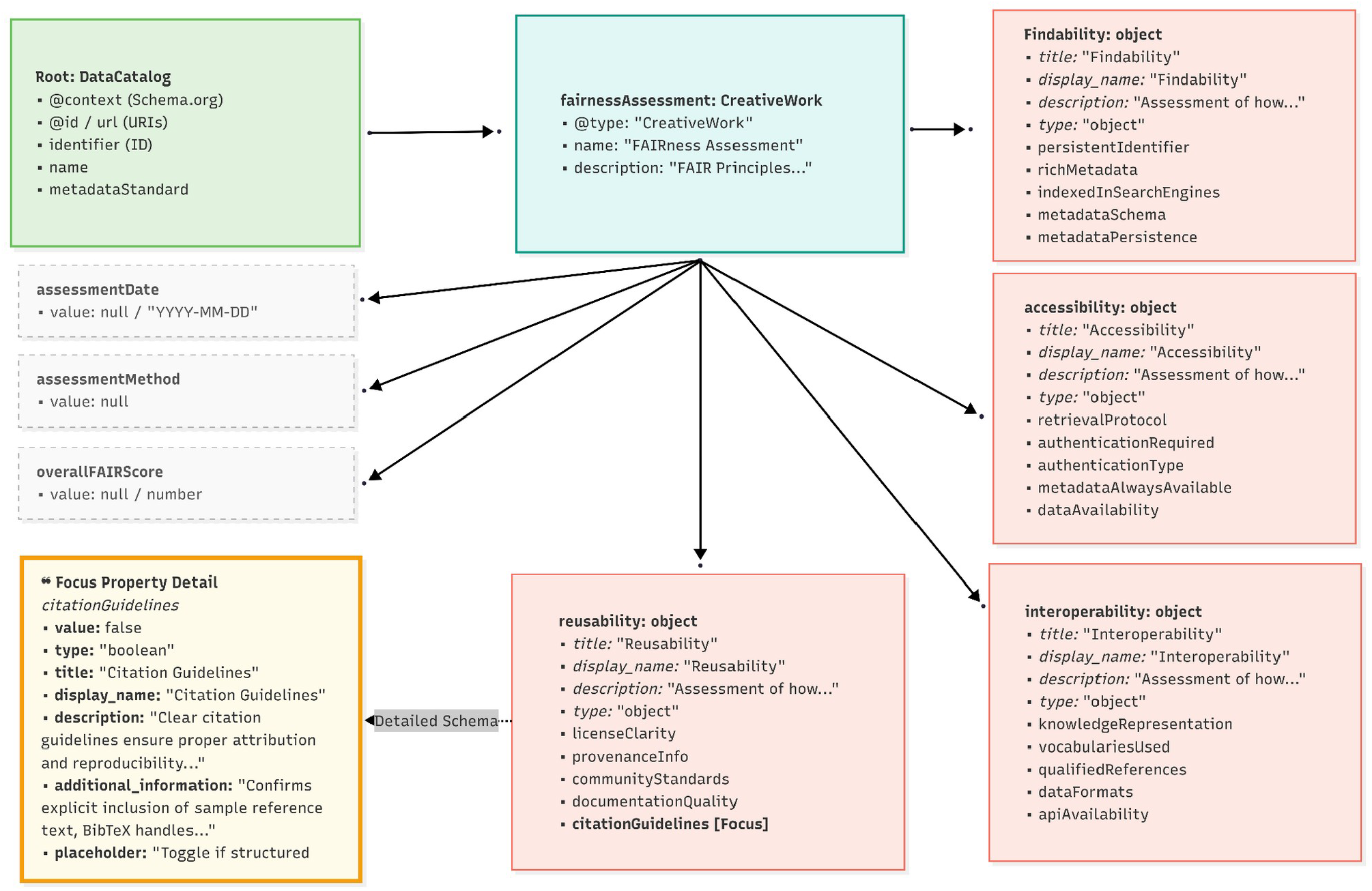
Machine-readable data structure architecture of the API payload. The entity inherits from the Schema.org @type: “DataCatalog” root tier, branching down into an isolated evaluation wrapper (fairnessAssessment). The framework represents the four core FAIR pillars not as flat keys, but as complex objects encapsulating structural metadata (title, display_name, description, type) alongside individual evaluation properties. The focused callout (bottom left) illustrates the comprehensive internal metadata profile allocated to a single primitive criterion (citationGuidelines). The image is generated using https://mermaid.js.org.

#### 4. Native Software Package Distribution

To lower the technical barrier to entry for third-party client applications, institutional portals, and external life-science platforms, the compiled repository inventory and its multi-dimensional FAIR attributes have been completely modularised. The entire validated property catalogue is distributed as an open-source, strongly typed software dependency via the public npm registry under the identifier rf-rdis (https://www.npmjs.com/package/rf-rdis). This approach allows web developers and automated workflows across the broader NFDI and EOSC ecosystems to natively install, filter, and embed the verified agrosystem repository index directly into their own local runtime environments and CI/CD pipelines without incurring network latency or REST API overhead.

The package is built with TypeScript, supplying native typing interfaces (Rdi) to support compile-time safety and IDE autocompletion for downstream integrations. Rather than relying on external web requests, the package bundles all RDI evaluation JSON files directly in its distribution. It exposes a minimal, highly performant API surface for querying this data in-memory:

The following snippet demonstrates how a downstream client application or automated pipeline installs the library and programmatically evaluates repository characteristics locally.

Installation command:

npm install rf-rdis

In-memory metadata querying using the rf-rdis library:

import {getRdiById, getFairnessById, searchRdis} from ‘rf-rdis’;

// 1. Resolve a specific repository locally using its PID const repository = getRdiById(‘RFId001202604272’);

console.log(‘Loaded repository: ${repository?.generalInfo?.title}’);

// 2. Fetch the corresponding structured FAIRness assessment object const fairnessPayload = getFairnessById(‘RFId001202604272’); const overallScore = fairnessPayload?.overallFAIRScore?.value; console.log(‘Evaluated overall FAIR Score: ${overallScore}%’);

// 3. Filter repositories by custom agronomic domain predicates offline const soilRepositories = searchRdis(rdi => rdi.generalInfo?.title?.toLowerCase().includes(‘soil’)

);

## Discussion

Our framework transitions FAIR assessment from a subjective, checklist-driven exercise into a standardized, data-driven service. By operationalizing these multi-dimensional metrics and publishing the live records via the FAIRagro Search Hub, we establish a transparent, citable “FAIRness benchmark”. This architecture incentivizes infrastructure operators to engage in targeted optimization, establishing a clear operational pathway that mirrors the broader community vision for structured “FAIRification” roadmaps (Jacobsen et al., 2020).

By deliberately avoiding a single, composite FAIR index, this methodology ensures that the nuance of repository performance is not lost in mathematical averages. This granular approach makes it immediately apparent where an infrastructure excels and where it lacks. For instance, an RDI might boast high accessibility through robust protocols for data retrieval but be nearly invisible due to poor Findability. Our multi-dimensional scoring approach addresses the common criticism that aggregate FAIR scores can mask specific technical deficiencies (Devaraju & Huber, 2021). For instance, by isolating “Interoperability” from “Findability,” we provide RDI managers with a clear ‘FAIRification’ roadmap for infrastructure optimization, such as improving metadata persistence or the use of controlled vocabularies, instead of guessing why their FAIRness index is low, or how to improve it.

A significant insight gained during the deployment of this framework is the current limitation of centralized, global repository directories. While registries like re3data provide a valuable baseline for high-level repository discovery, their underlying schemas are inherently optimized for generic, administrative tracking rather than deep technical evaluation. The presented pipeline addresses this limitation by formalizing a strict data-provenance hierarchy across re3data-native, inferred and novel properties. This architecture demonstrates how domain-specific consortia can build on top of global infrastructure without duplicating efforts, by programmatically inferring hidden technical capabilities and carving out a dedicated tier for novel, domain-specific indicators (e.g., agricultural ontology compliance, dataset provenance tracking). The developed methodology contributes to solving the classic conflict between automated scalability and data fidelity. The automated enrichment pipeline ensures the index scales efficiently across external infrastructures, while the novel attributes from the continuous interview process and validation loop guarantee that the highly specific, high-fidelity characteristics required by modern AI and big-data workflows are accurately captured.

Perhaps the most significant departure from traditional FAIR assessment frameworks is how this metadata is deployed. Historically, FAIR evaluations have been treated as static artifacts, PDF reports or isolated web dashboards that exist separate from actual scientific computing. The integration of this framework into the FAIRagro ecosystem completely reframes FAIR metadata as a live, runtime resource. By exposing the validated repository property dataset simultaneously through a human-centric visual UI, an automated REST API, and an open-source npm package (rf-rdis), the framework shifts the paradigm from passive compliance reporting to active infrastructure dependency. Third-party developers, NFDI search hubs, and client-side analysis tools can now programmatically pull the live JSON payloads to dynamically filter repositories based on real-time feature profiles. This allows users to establish highly customised, requirement-driven repository rankings based on their immediate programmatic needs, entirely bypassing rigid, predefined scoring monopolies.

Ultimately, this methodology does more than just evaluate repositories; it fosters a culture of technical accountability and transparent data stewardship across a historically fragmented agrosystems landscape. Because this framework aligns directly with the strategic mandates of the German National Research Data Infrastructure (NFDI) and the overarching European Open Science Cloud (EOSC) vision (Dillo et al. 2025), it significantly lowers the barrier to entry for other life science consortia. Instead of developing bespoke validation frameworks from scratch, neighboring disciplines can ingest and replicate this scripted pipeline to benchmark their own decentralized infrastructures. This collaborative, open-source model drives the sustainable adoption of FAIR standards, ensuring that federated agricultural research remains highly reproducible, machine-actionable, and globally integrated.

## Conclusion and Outlook

This methodology establishes a scalable and reproducible framework for multi-dimensional infrastructure assessment within the German National Research Data Infrastructure (NFDI). By operationalizing the abstract criteria of the RDA FAIR Data Maturity Indicators into twenty core repository-level metrics, this initiative delivers a technical baseline that respects the operational autonomy of decentralized nodes while systematically driving cross-disciplinary interoperability.

The dual-source ingestion pipeline, combining direct operator interviews with automated metadata harvesting, successfully demonstrates how domain-specific consortia can enrich global registries like re3data without duplicating curation efforts. Crucially, by rejecting single, composite scores in favor of independent, pillar-specific ratings, the framework ensures that a repository’s distinct architectural strengths and critical technical bottlenecks remain completely transparent to both users and infrastructure managers.

Furthermore, the live deployment of this property catalogue through the FAIRagro Search Hub prototype transitions FAIR assessment from a passive, retrospective auditing exercise into an active runtime software utility. Exposing validated infrastructure metadata simultaneously via interactive visual dashboards, a dedicated RESTful API, and an open-source software dependency (rf-rdis via npm) provides the broader life-science community with the machine-actionable tools necessary to natively integrate data stewardship indicators directly into automated developer workflows, federated search engines, and local data-management pipelines.

Future work will focus on expanding the programmatic parsing capabilities of the automation pipeline to further reduce reliance on manual survey instruments and curation files. By refining the text-mining and heuristic rules within the ingestion module, future iterations will aim to automatically extract complex technical indicators, such as authentication protocols and controlled vocabulary compliance, directly from public repository documentation and endpoint landing pages. Additionally, efforts will be directed toward advocating for the structural integration of our newly defined, technical properties, not indexed by registries, into upcoming iterations of global metadata standards.

Ultimately, this hybrid approach proves that combining transparent, data-driven scoring with the consensus evaluations of an expert editorial board serves as the most effective mechanism for fostering technical accountability and accelerating the sustainable adoption of FAIR standards across decentralized research landscapes.

## Author Contributions

### Ataul Haleem (AH)

Responsible lead for conceptualization, methodology, investigation, data acquisition and curation (API integration, automated harvesting, and data analysis and visualization), software (architecture, API design, and full-stack interactive prototype implementation) and writing the original draft. AH also conducted the interviews with RDI managers for Repository FAIRness assessment.

### Elena Rey Mazón (EM) & Marcus Schmidt (MS)

Involved in data curation, validation (Editorial Board expert peer-review), and review and editing.

### Jagadeeshwar Reddy Etukala (JE)

Involved in data acquisition.

### Jascha Jung (JJ), Daniel Martini (DM), & Daniel Arend (DA)

Supported the overall inventory technical alignment, writing, reviewing, and editing.

### Charlotte Neidiger & Robert Ulrich

Supported data complementation of new agrosystem RDIs within the re3data registry to facilitate clean metadata harvesting. Additionally involved in data curation, validation, review and editing.

### Björn Usadel (BU)

Measure lead (M4.3), funding acquisition, review and editing.

### Matthias Lange (ML)

Responsible for funding acquisition, project administration, and review and editing. ML actively provided critical feedback and was involved in the development of the methodology and the infrastructure.

## Acknowledgments

The authors would like to express their sincere gratitude to the RDI managers from the participating infrastructures who contributed their valuable time and insights during the initial phase of this ongoing integration:

- BacDive
- BonaRes Repository
- CCDC (Cambridge Crystallographic Data Centre)
- DWD OpenData (Deutscher Wetterdienst)
- Edaphobase
- EDI
- ioChem-BD
- OpenAgrar
- PANGAEA
- PhenoRoam
- Thünen Atlas
- ZB MED LifeData
- Zenodo
- EUDAT B2SHARE

---

**Supplementary Figure 1:**
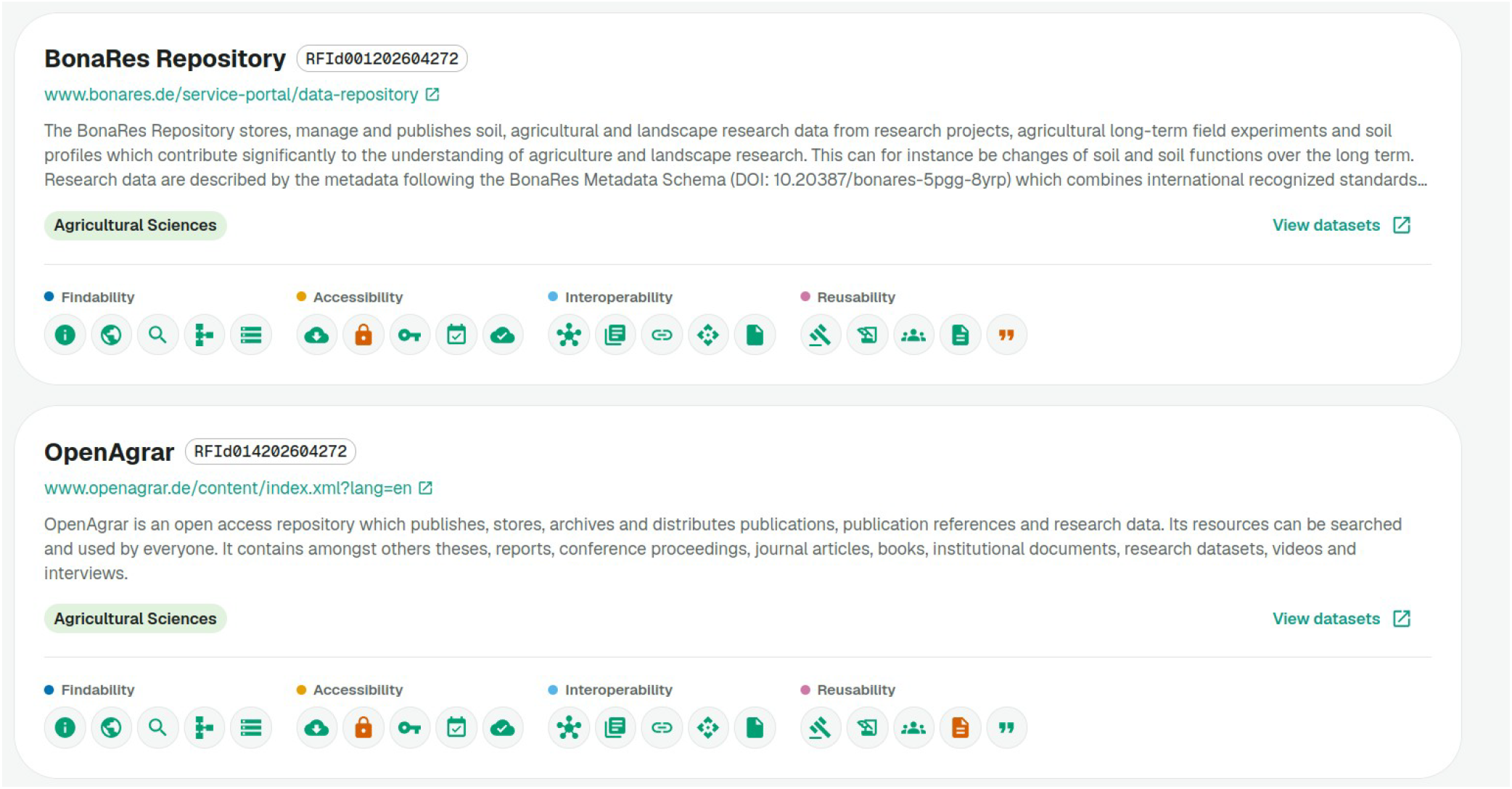
A comparative view resulting from a search term ‘agricultural’.

**Supplementary Figure 2:**
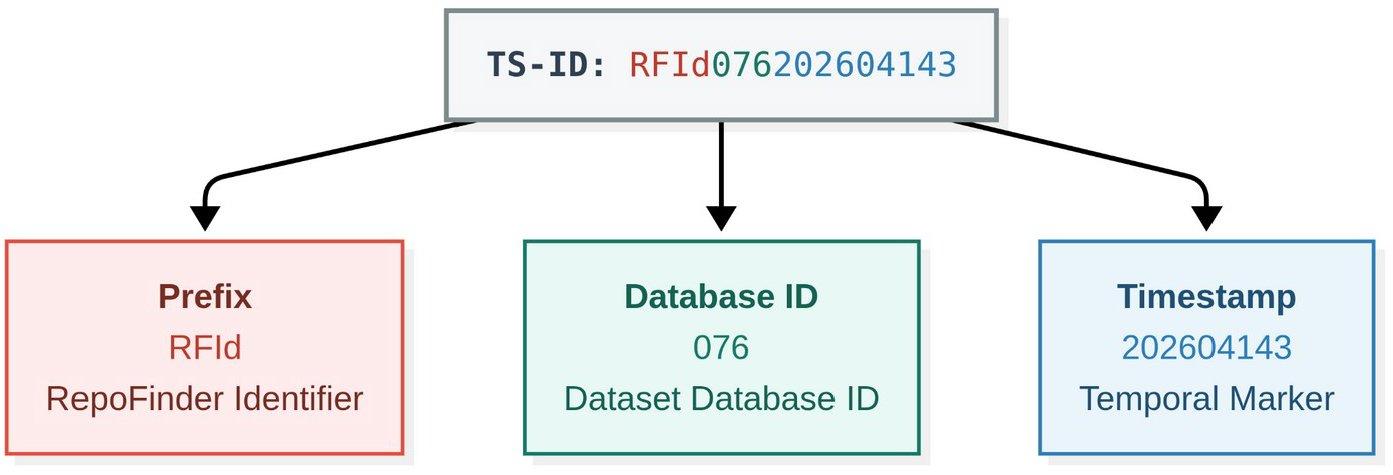
Anatomy of a RepoFinder identifier (RFId). RFIds are assigned once when a repository is registered in the inventory and are never regenerated. The dummy example RFId076202604143 decomposes into a fixed namespace prefix (RFId), a zero-padded database index (076) that ensures uniqueness within the inventory, and a registration-date marker (202604143, the date 2026-04-14 followed by a weekday digit) that records when the entry was created. Diagram generated with Mermaid (https://mermaid.js.org).

**Supplementary Table 1:**
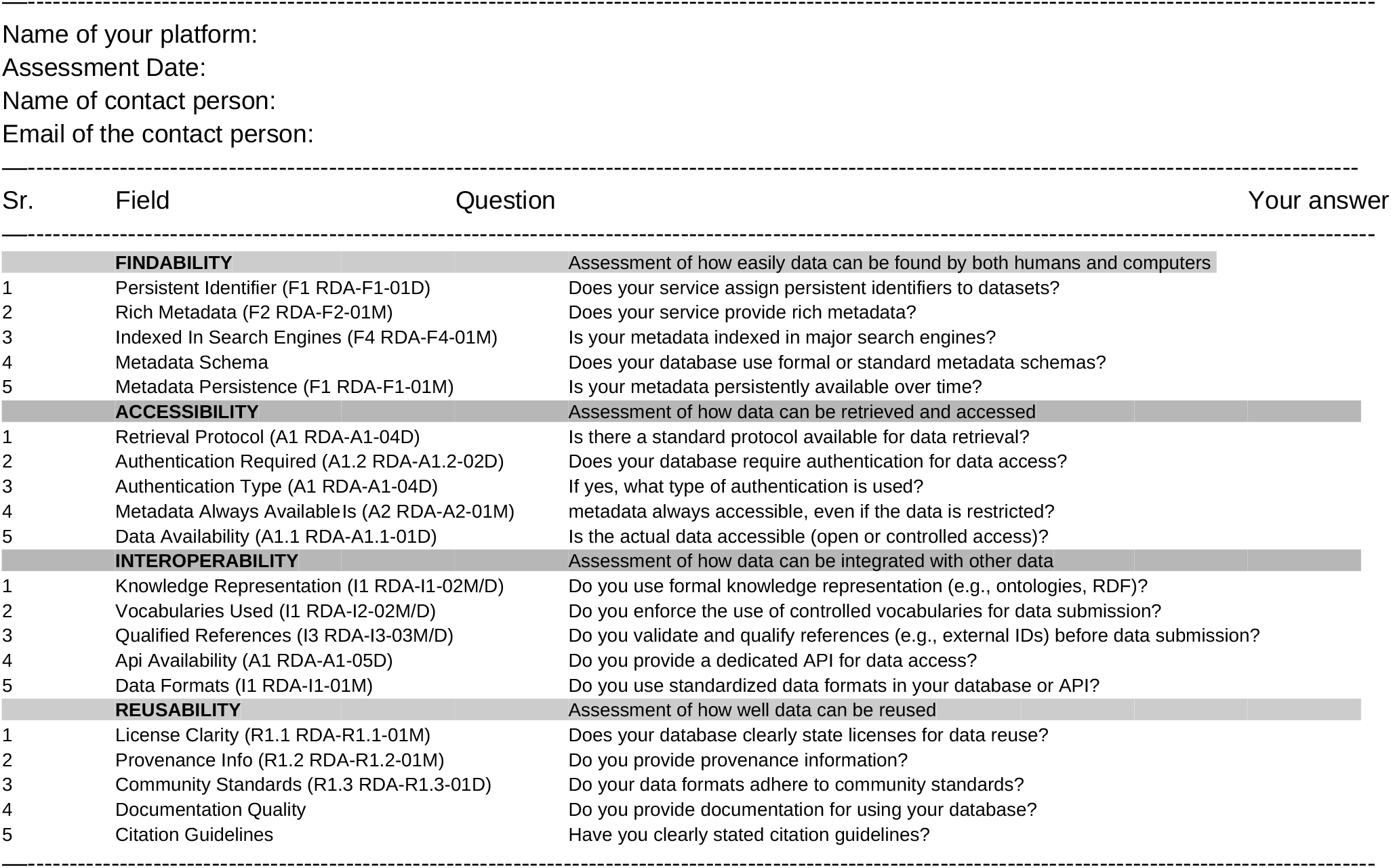
Interview questionnaire for RDI managers.

## References

Azevedo, L. G., Banaggia, G., Tesolin, J., & Cerqueira, R. (2025). Analysis of automated tools for FAIRness evaluation: A literature perspective. In Lecture Notes in Computer Science (pp. 149–166). Springer Nature Switzerland.

Candela, L., Mangione, D., & Pavone, G. (2024). The FAIR assessment conundrum: Reflections on tools and metrics. Data Science Journal, 23(1), 33.

Curdt, C., Dierkes, J., & Kloppenburg, S. (2022). RDM in a decentralised university ecosystem —A case study of the university of cologne. Data Science Journal, 21(20). 10.5334/dsj-2022-020

Devaraju, A., & Huber, R. (2020). F-UJI - an automated FAIR data assessment tool. Zenodo. 10.5281/ZENODO.4063720

Devaraju, A., & Huber, R. (2021). An automated solution for measuring the progress toward FAIR research data. Patterns (New York, N.Y.), 2(11), 100370.

Diepenbroek, M., Glöckner, F. O., Grobe, P., Güntsch, A., Huber, R., König-Ries, B., Kostadinov, I., Nieschulze, J., Seeger, B., Tolksdorf, R., & Triebel, D. (2014). Towards an integrated biodiversity and ecological research data management and archiving platform: the German federation for the curation of biological data (GFBio). Informatik 2014, 1711–1721.

Diepenbroek, M., Kostadinov, I., Seeger, B., Glöckner, F. O., Dieckmann, M. A., Goesmann, A., Ebert, B., Schimmler, S., & Sure-Vetter, Y. (2023). Towards a Research Data Commons in the German National Research Data Infrastructure NFDI: Vision, governance, architecture. Proceedings of the Conference on Research Data Infrastructure, 1. 10.52825/cordi.v1i.355

Dillo, I., Verburg, M., Davidson, J., Jonquet, C., & Aubin, S. (2025). Expanding FAIR solutions across EOSC. https://hal.science/hal-05085825/

Gaignard, A., Rosnet, T., De Lamotte, F., Lefort, V., & Devignes, M.-D. (2023). FAIR-Checker: supporting digital resource findability and reuse with Knowledge Graphs and Semantic Web standards. Journal of Biomedical Semantics, 14(1), 7.

Jacobsen, A., Kaliyaperumal, R., da Silva Santos, L. O. B., Mons, B., Schultes, E., Roos, M., & Thompson, M. (2020). A generic workflow for the data FAIRification process. Data Intelligence, 2(1-2), 56–65.

Maemura, E. (2021). Data here and there: Studying web archives research infrastructures in Danish and Canadian settings. https://utoronto.scholaris.ca/bitstreams/995f62ff-56dd-4ac9-b982-d8a11b7369b1/download

Mons, B., Neylon, C., Velterop, J., Dumontier, M., da Silva Santos, L. O. B., & Wilkinson, M. D. (2017). Cloudy, increasingly FAIR; revisiting the FAIR Data guiding principles for the European Open Science Cloud. Information Services & Use, 37(1), 49–56.

RDA FAIR Data Maturity Model Working Group, B. (2020). FAIR Data Maturity Model: specification and guidelines. Research Data Alliance. 10.15497/rda00050

Schlindwein, S. L. (2025). Breaking down disciplinary silos? In search of interdisciplinary integration in postgraduate agroecosystems research. Kybernetes. The International Journal of Cybernetics, Systems and Management Sciences, 54(11), 6507–6518.

Stockhause, M., & Lautenschlager, M. (2017). CMIP6 data citation of evolving data. Data Science Journal, 16, 30.

Wilkinson, M. D., Dumontier, M., Aalbersberg, I. J. J., Appleton, G., Axton, M., Baak, A., Blomberg, N., Boiten, J.-W., da Silva Santos, L. B., Bourne, P. E., Bouwman, J., Brookes, A. J., Clark, T., Crosas, M., Dillo, I., Dumon, O., Edmunds, S., Evelo, C. T., Finkers, R., … Mons, B. (2016). The FAIR Guiding Principles for scientific data management and stewardship. Scientific Data, 3(1), 160018.

Zänkert, S., Manske, A., Miller, B., & Fluck, J. (2025). Base4NFDI: Fostering A cross-disciplinary service landscape for the German National Research Data Infrastructure. International Journal of Digital Curation, 19(1), 7.

